# WISP1 drives a mechanically active immune modulatory and proliferative cardiac myofibroblast state

**DOI:** 10.64898/2026.02.17.706476

**Authors:** Sharon Parkins, Sarah R. Anthony, Teagan K. Goldsworthy, Akanksha Nigam, Neva C. Schehl, Robert M. Jaggers, Daniel A. Kasprovic, Lisa C. Green, Onur Kanisicak, Michael Tranter

## Abstract

Pathological cardiac remodeling is driven by the proliferation and differentiation of resident fibroblasts into active myofibroblasts and results in excessive extracellular matrix (ECM) deposition and tissue stiffening. Expression of the matricellular protein WISP1 has previously been shown to be increased with cardiac fibrosis and promote myofibroblast activity, but the mechanisms by which this occurs remain unknown. Primary cardiac fibroblasts were isolated from adult mouse hearts and treated with recombinant WISP1 or TGFβ1 both alone and in combination to determine the functional role of the matricellular protein WISP1 in driving cardiac myofibroblast activity. WISP1 significantly increased alpha-smooth muscle actin and collagen type I expression, total collagen secretion, collagen gel contractility, and wound healing equally in fibroblasts from both male and female mice. However, WISP1 alone failed to induce expression of periostin, a hallmark myofibroblast marker, suggesting the resulting WISP1-dependent cell phenotype is unique and/or acting through non-canonical pathways. Indeed, inhibition of P38 MAPK completely ablated the WISP1-dependent increase in *α*SMA and collagen expression, while having little to no impact on TGFβ1-dependent expression of myofibroblast marker genes. We next employed a multi-omics approach to define the functional impact of WISP1 on fibroblast cell-state within the transcriptome, cytosolic, and secreted ECM proteome. RNA-seq results show that WISP1 broadly promotes the expression of proliferative and immune modulatory genes at the transcriptomic level, while having very little impact on traditional myofibroblast and ECM modifying gene expression programs. At the proteome level, WISP1 was again a much weaker mediator of traditional myofibroblast and ECM proteins. However, in agreement with RNA-seq data, we observed a strong WISP1-dependent enrichment for proliferation-associated proteins in the cytosolic proteome and inflammation-associated proteins in the ECM proteome. Interestingly, WISP1 also showed a context-dependent response with TGFβ1, suggesting a more complex and yet to be elucidated signaling interaction between these independent mediators of myofibroblast activity. In conclusion, our data suggests that WISP1 promotes a unique proliferative and immune-modulatory myofibroblast phenotype.

**Highlights:**

- WISP1 is sufficient to drive myofibroblast *α*SMA and collagen expression and ECM deposition
- WISP1 promotes canonical myofibroblast contractility and wound healing activity
- WISP1 mediates myofibroblast activity via a non-canonical, P38 MAPK-dependent signaling pathway
- Multi-omics analysis of WISP1-dependent RNA and protein expression show that WISP promotes a proliferative and immune modulatory myofibroblast phenotype

## 1. Introduction

Pathological cardiac remodeling characterized by fibrosis is a hallmark of heart failure (HF) and a primary determinant of ventricular dysfunction [1–4]. While structural myocardial remodeling acts as an essential reparative mechanism to preserve mechanical integrity following injuries such as myocardial infarction (MI) [5–7], the persistent and unchecked accumulation of extracellular matrix (ECM) proteins, or fibrosis, often becomes maladaptive [3, 8]. This excessive ECM remodeling stiffens the ventricular wall, impairs diastolic and systolic compliance, and disrupts electrical conduction [9, 10, 11]. Despite advances in neurohormonal blockade using angiotensin-converting enzyme inhibitors, β-blockers, and mineralocorticoid receptor antagonists, these interventions only delay disease progression. Effective therapies that directly target the cellular effectors of fibrotic tissue remodeling remain elusive [8, 9, 12, 13].

The cardiac fibroblast is the principal cell type responsible for maintaining ECM homeostasis in the myocardium [14]. Under physiological conditions, fibroblasts are quiescent; however, in response to ischemic injury or mechanical stress, they undergo a defined differentiation program into proliferation-competent, α-smooth muscle actin (α-SMA) positive myofibroblasts [15,16]. These activated myofibroblasts serve as the primary source of interstitial collagen types I and III, fibronectin, and matrix metalloproteinases (MMPs) required for scar formation [17, 18]. While the initial recruitment of myofibroblasts consolidates the infarcted region and prevents rupture, their persistence in the chronic phase drives interstitial fibrosis in the non-infarcted remote myocardium [16]. Consequently, delineating the molecular mechanisms that mediate and sustain myofibroblast activity is critical for developing targeted anti-fibrotic strategies.

Transforming growth factor-β1 (TGFβ1) is the canonical mediator of fibroblast activation and fibrosis [19–22]. The CCN family of matricellular proteins have also been shown to serve as vital mediators of cell adhesion, migration, and cell-matrix communication [23]. Among these, Wnt1-inducible signaling pathway protein-1 (WISP1/*CCN4*) has emerged as a new mediator of fibroblast activity across multiple organ systems. WISP1 is markedly upregulated in idiopathic pulmonary fibrosis (IPF), where it functions as a key downstream effector of TGFβ1 signaling to drive fibroblast proliferation and matrix synthesis [24] and drives stage-dependent fibrogenesis in human dermal and lung fibroblasts [25,26]. WISP1 expression has also been shown to positively correlate with fibrosis in human liver in NASH patients, and WISP1 inhibition prevented the progression, but not onset, of hepatic fibrosis. WISP1 expression in the healthy adult heart is low, but is markedly increased following MI specifically within the non-ischemic myocardium [27]. Limited existing work regarding the role of WISP1 on cardiac fibroblasts suggests that WISP1 promotes proliferation, migration, SMA expression, and collagen expression and processing [28–29]. Despite these observations, the precise signaling networks through which WISP1 modulates fibroblast function, how WISP1 integrates with TGFβ1 signaling, and the functional state of a WISP1-activated fibroblast remain poorly understood.

Here, we sought to determine the role WISP1 plays in regulating cardiac fibroblast activation and extracellular matrix remodeling, both independently and in coordination with TGFβ1. To define the WISP1-dependent myofibroblast cell state, we used a multi-omics approach that integrated transcriptomics with the cytosolic and ECM proteome. We found that while WISP1 induces the expression of some canonical myofibroblast markers, the predominate effect of WISP1 drives a proliferative and immune modulatory cell state.

## 2. Methods

### 2.1. Animal Husbandry and Isolation, Culture, and Treatment of Primary Murine Cardiac Fibroblasts

All mouse studies were approved by the University of Cincinnati and The Ohio State University Institutional Animal Care and Use Committee (IACUC) under protocol numbers 22-11-15-02 and 2023A00000092, respectively, and comply with the ARRIVE guidelines. All animal experiments were performed in accordance with these guidelines.

Primary cardiac fibroblasts (CFs) were isolated from left ventricular tissue of 8–12-week-old C57BL/6 mice (Jackson Laboratory, stock #: 000664) as previously described [30]. Briefly, excised ventricles were minced and digested in HBSS containing Collagenase IV (Worthington, LS004189) and Dispase II (Sigma-Aldrich, D4693) at 37 °C on a rocker for 45–60 min, with gentle pipetting every 15 min. The digest was filtered through a 40 µm cell strainer (Corning, 431750) and centrifuged at 300 × g for 10 min. The pellet was resuspended in DMEM with 10% FBS (Gibco) and pre-plated for 45 min to enrich fibroblasts by plastic adherence. Non-adherent cells were removed, and adherent CFs were expanded to ∼70% confluence.

For treatments, CFs were seeded in 12- or 24-well plates in DMEM + 10% FBS and treated for 72 h with recombinant human TGFβ1 (10 ng/mL; R&D Systems, 7754-BH) and/or recombinant murine WISP1 (500 ng/mL; R&D Systems, 1680-WS-050). Where indicated, CFs were serum-starved in Opti-MEM (Gibco) and pre-treated with SB203580 (10 µM; Tocris Bioscience) for 30 min before addition of growth factors. DMSO (Sigma-Aldrich) at equivalent concentration served as vehicle control.

### 2.2. RNA Isolation and Quantitative PCR

Total RNA was extracted using the NucleoSpin RNA kit (Macherey-Nagel, 740955.250) per manufacturer’s instructions. First-strand cDNA was synthesized with iScript Reverse Transcription Supermix (Bio-Rad, 1708841). qPCR was performed on a Bio-Rad CFX96 using iTaq Universal SYBR Green Supermix (Bio-Rad, 1725121). Relative gene expression was calculated by the ΔΔCt method with 18S as reference. Primer sequences used were as follows:

18S: F 5′-AGTCCCTGCCCTTTGTACACA-3′; R 5′-CCGAGGGCCTCACTAAACC-3′

Postn: F 5′-GGTGGCGATGGTCACTTATT-3′; R 5′-GTCAGTGTGGTGGCTCTTAC-3′

Col1a1: F 5′-CTGGCGGTTCAGGTCCAA-3′; R 5′-GCCTCGGTGTCCCTTCATT-3′

WISP1: F 5′-CACCACTAGAGGAAACGACTAC-3′; R 5′-TCACAGCCATCTGTGATTAGG-3′

### 2.3. Flow Cytometry

To determine the baseline binding capacity of WISP1 to cardiac fibroblasts, primary murine cardiac fibroblasts were isolated and cultured as previously described. Recombinant WISP1 was conjugated to PE using the Lightning-Link® PE/R-Phycoerythrin Conjugation Kit (Abcam, ab102918) according to the manufacturer’s instructions. A saturation binding curve was generated using PE-labeled recombinant WISP1 (rWISP1-PE).

Briefly, harvested cells (1.0–1.5 x 10⁵ cells/well) were detached using Accutase (ThermoFisher, 00-4555-56) and incubated with murine Fc Block (BioLegend, 156604, 0.25ug/100ul) for 15 minutes at 4°C. Cells were then subjected to an 8-point serial dilution of rWISP1-PE in FACS buffer made of DPBS (Gibco, 14190-136) with 0.09% Sodium Azide and 1% BSA, ranging from 0.0005 µg/mL to 5.0 µg/mL. Following a 1-hour incubation at 4°C, cells were washed twice with FACS buffer and stained 7-AAD (ThermoFisher, A1310, 1:2000) at 4°C for 5 minutes to exclude dead cells from the final analysis. Data were acquired on a Cytek Bioscience Aurora Spectral Analyzer with a five laser configuration of: 355, 405, 488, 561, 640 nm, and 64 detection channels. The data were analyzed using FlowJo analysis software (BD, V10.10). Unstained cells were used for autofluorescence extraction, along with a single stained rWISP1-PE reference control to define negative and positive gates. The percentage of PE-positive cells was used to calculate the dissociation constant (Kd) and maximum binding (Bmax) using non-linear regression analysis.

### 2.4. Protein Isolation and Western Blotting

CFs treated for 72 h were lysed in RIPA buffer supplemented with 0.5 mM DTT, 0.2 mM sodium orthovanadate, and Halt Protease & Phosphatase Inhibitor Cocktail (Thermo Fisher, 78443). Four micrograms of protein per lane were resolved on 10% TGX Stain-Free gels (Bio-Rad, 4568035), transferred to nitrocellulose membranes, and blocked in 5% BSA (MilliporeSigma, A7906) in Tween-Tris Buffered Saline (T-TBS) for 1 h at room temperature. Membranes were incubated overnight at 4 °C with primary antibodies (1:1,000 in 5% BSA): Periostin (Novus Biologicals, NBP1-30042), α-SMA (Cell Signaling, D4K9N), Col1a1 (Cell Signaling, E8F47), Fibronectin (Cell Signaling, E7F5X), and GAPDH (Cell Signaling, 14C10). HRP-conjugated anti-rabbit secondary antibody (Bio-Rad, 1721019; 1:10,000) was applied for 1 h at room temperature. Signal was captured using chemiluminescent substrate (Thermo, 34580) on a ChemiDoc Imaging System and band density was quantified using ImageJ and normalized to GAPDH.

### 2.5. Immunofluorescence

CFs cultured in glass-bottom 24-well plates were fixed in 4% PFA for 10 min, permeabilized in 0.1% Triton X-100 for 10 min, and blocked in 2% BSA (Sigma-Aldrich, A3059) for 1 h at room temperature. Primary antibodies (1:250 in 2% BSA) against vimentin (Abcam, ab8978) α-SMA (Abcam, ab7817) and fibronectin (ABCAM EPR19241-46] were applied for 2 h at room temperature or overnight at 4°C. Next, Alexa Fluor 488 anti-mouse secondary antibody (Abcam, ab150113; 1:250) and/or Alexa Fluor 647 anti-rabbit secondary antibosy (Abcam, ab150079) was applied for 1 h at room temperature. Nuclei were counterstained with ProLong Diamond Antifade Mountant with DAPI (Invitrogen, P36962). Images were acquired on a BioTek Cytation 10 and analyzed in ImageJ.

### 2.6. Wound Closure Assay

CFs were plated in 96-well plates and grown to ∼75% confluence. Scratches were created using a BioTek AutoScratch tool, wells were washed with PBS, and treatments were applied as described in Section 2.1. Time-lapse imaging was performed on a BioTek Cytation 10, and wound closure was quantified using the *MRI_Wound_Healing_Tool* macro in ImageJ.

### 2.7. Collagen Gel Contraction Assay

Collage gel contraction assays were conducted using the CytoSelect™ 48-Well Floating Matrix Kit (Cell Biolabs, CBA-201). Collagen gels (1 mg/mL) were seeded with 1.5 × 10^5 CFs per well and polymerized at 37 °C for 1 h per manufacturer’s instructions. Treatments were added, gels were released, and images were captured at 0, 6, 12, and 24 h. Gel surface area was measured in ImageJ.

### 2.8. Total Collagen Production

After 72 h of treatment, CFs were fixed in ice-cold methanol overnight at –20 °C. Wells were rinsed with PBS, stained with Picrosirius Solution B (Polysciences, 24901B) for 1 h at room temperature, washed twice with 0.1 N HCl, and eluted in 0.1 M NaOH for 1 h at room temperature as previously described [30]. Absorbance at 540 nm was measured and imaged on a Cytation 10.

### 2.9. RNA-sequencing and bioinformatics analysis

Bulk RNA sequencing was performed by The Ohio State University Comprehensive Cancer Center Genomics Shared Resources Core at read depth of ∼40M paired end reads per sample. Reads were then mapped and quantified using Rsubead (v2.18.0) in R (v4.5.1). Raw transcript-level counts and sample metadata were imported into and processed using DESeq2 (v1.44.0). A multifactor linear model was used to quantify TGF-β1 main effect, Wisp main effect, and Wisp × TGF-β1 interaction. Gene identifiers (Ensembl IDs) were mapped to murine gene symbols using org.Mm.eg.db (v3.19.1). Variance-stabilized expression values were computed using VST (variance stabilizing transformation) and ranked gene lists were generated for each contrast prior to performing Gene Set Enrichment Analysis (GSEA) using MSigDB Hallmark gene sets.

### 2.10. Cytosolic and ECM Protein Isolation and Mass Spectrometry

CFs were plated and treated as described in Section 2.1. To harvest cytosolic and ECM proteins, cells were cultured for six days (three days in DMEM + 10% FBS to reach confluence, followed by three days under treatment conditions). Media was removed and cells were washed twice with PBS. Cells were lysed in situ with RIPA buffer supplemented with 0.5 mM DTT, 0.2 mM sodium orthovanadate, and Halt Protease & Phosphatase Inhibitor Cocktail (Thermo Fisher, 78443). Without disturbing the deposited ECM, the lysate was collected and centrifuged at 14,000 × g for 15 min at 4 °C. The supernatant (soluble proteome fraction) was transferred to new tubes, quantified by BCA assay (Pierce, 23225), and stored at –80 °C until mass-spectrometry processing. Following soluble protein harvest, the remaining ECM was gently washed with PBS and decellularized in 0.25% Triton X-100/0.25% sodium deoxycholate for 24 h at 37 °C. Wells were then rinsed three times with PBS and the decellularized ECM was solubilized in 2.5% SDS/50 mM Tris (pH 6.8) and manually dissociated from wells by scraping.

Samples were analyzed by nanoLC-MS/MS (Orbitrap Eclipse) at The Ohio State University CCIC Mass Spectometry and Proteomics Facility. Data were processed in Proteome Discoverer 2.4 with the Sequest HT algorithm for label-free quantitation without normalization. Protein-level abundances (global/soluble and ECM fractions) were imported into R (v4.5.1) for downstream analysis. Differential protein abundance was modeled using limma (v3.66.0) with a factorial design to estimate the TGFβ1 main effect, Wisp main effect, and Wisp × TGFβ1 interaction. To account for repeated measures arising from matched biological donors across treatment conditions, within-mouse correlation was estimated using duplicateCorrelation, and linear models were fit using lmFit with mouse as a blocking factor, followed by empirical Bayes moderation.

To quantify compartment-specific regulation, an ECM-enriched protein matrix was generated by subtracting global (soluble) protein abundances from matched ECM abundances for proteins detected in both fractions. Ranked protein lists from each contrast were used for pathway-level analysis by FGSEA using MSigDB Hallmark gene sets, analogous to the RNA-seq workflow.

### 2.11. Statistical Analysis

Quantitative results are presented as mean ± SEM. Statistical differences between groups were determined using Student’s t-tests or analysis of variance (ANOVA), as appropriate. All analyses were performed using GraphPad Prism (version 10), with p < 0.05 defining statistical significance.

## 3. Results

### 3.1. WISP1 is induced by TGFβ1 and is sufficient to drive myofibroblast *α*SMA and collagen expression and ECM deposition

To characterize the expression kinetics of WISP1 during fibroblast activation, primary mouse cardiac fibroblasts were treated with TGFβ1 (10 ng/mL) followed by assessment of *Wisp1* mRNA expression at 24, 48, and 72 horus post-treatment. Results show a progressive increase of *Wisp1* expression that reached statistical significance at 48 hours and peaked at 72 hours relative to vehicle controls (Fig. 1A).

**Figure 1.**
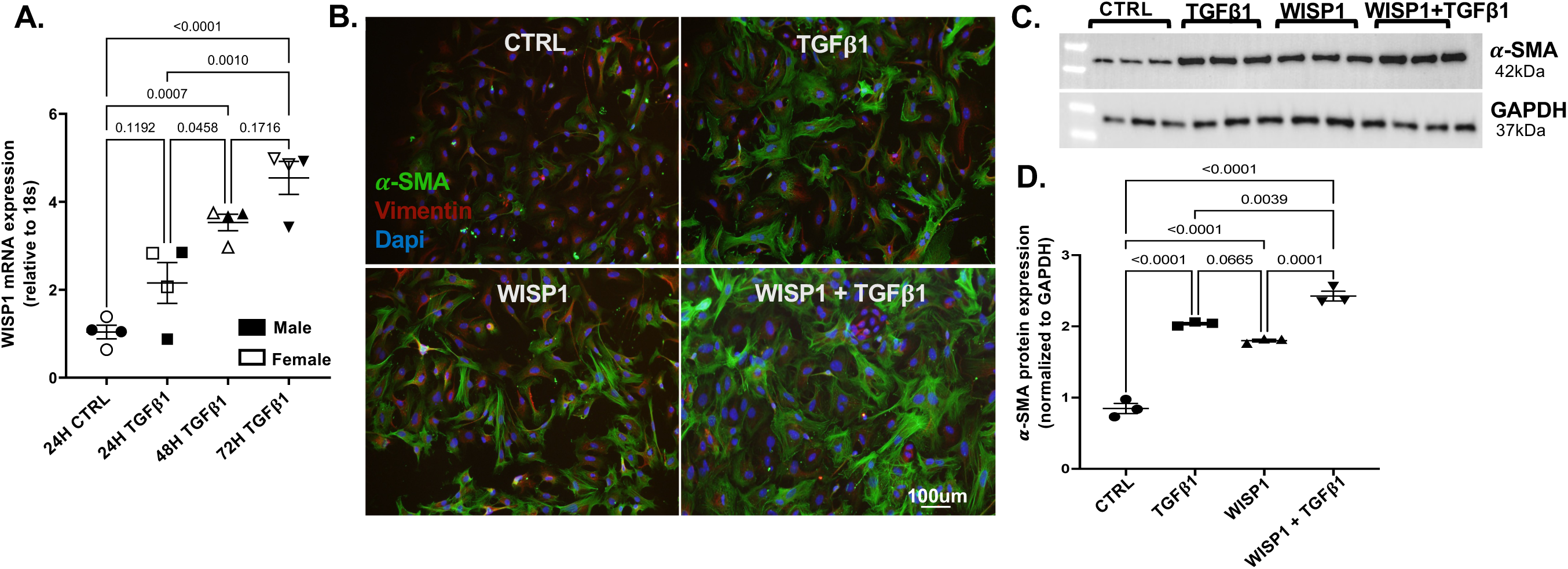
WISP1 is induced by TGFβ1 and is sufficient to drive α-SMA expression in cardiac fibroblasts. **(A)**Wisp1 mRNA expression in primary adult mouse cardiac fibroblasts following TGFβ1 (10 ng/mL) treatment for 24–72 h. **(B)** Representative immunofluorescence images of α-SMA (green), vimentin (red), and DAPI (blue) after 72 h treatment with vehicle (Ctrl), WISP1 (500 ng/mL), TGFβ1 (10 ng/mL), or WISP1 + TGFβ1. **(C, D)** Representative Western blot **(C)** and densitometric quantification **(D)** of α-SMA normalized to GAPDH. Data are mean ± SEM from ≥3 independent isolations. Statistical analysis by one-way ANOVA with Tukey’s post hoc test; *p < 0.05. Data are mean ± SEM from ≥3 independent isolations. Statistical analysis by one-way ANOVA with Tukey’s post hoc test; *p < 0.05.

We next assessed whether WISP1 alone is sufficient to drive the myofibroblast phenotype. To determine the optimal WISP1 dosing, we first measured WISP1 binding to cardiac fibroblasts using flow cytometry and observed an estimated WISP1 binding saturation at approximately 500 ng/mL (Fig. S1), consistent with previously published dosing strategy [28]. Fibroblasts were treated with TGFβ1 (10 ng/mL), WISP1 (500 ng/mL), or a combination of both for 72 hours. Immunofluorescence and Western blotting revealed that, similar to TGFβ1, WISP1 alone significantly increased the expression of α-SMA compared to controls (Fig. 1B-D). Interestingly, WISP1 and TGFβ1 together resulted in a further increase in α-SMA compared to either treatment alone.

We next determined the effect of WISP1 on ECM production by measuring total soluble collagen secretion using a picrosirius red assay. Treatment with WISP1 alone induced a significant increase in total collagen production compared to vehicle controls (Fig. 2A-B) as well as *Col1a1* mRNA (Fig. 2C) and collagen I protein expression (Fig. 2D-E). In contrast to α-SMA expression, WISP1 did not further potentiate TGFβ1-induced collagen production. Importantly, our results are consistent across cells isolated from both male and female mice, suggesting no sex difference in WISP1 or TGFβ1 regulated expression of α-SMA or collagen in isolated cells.

**Figure 2.**
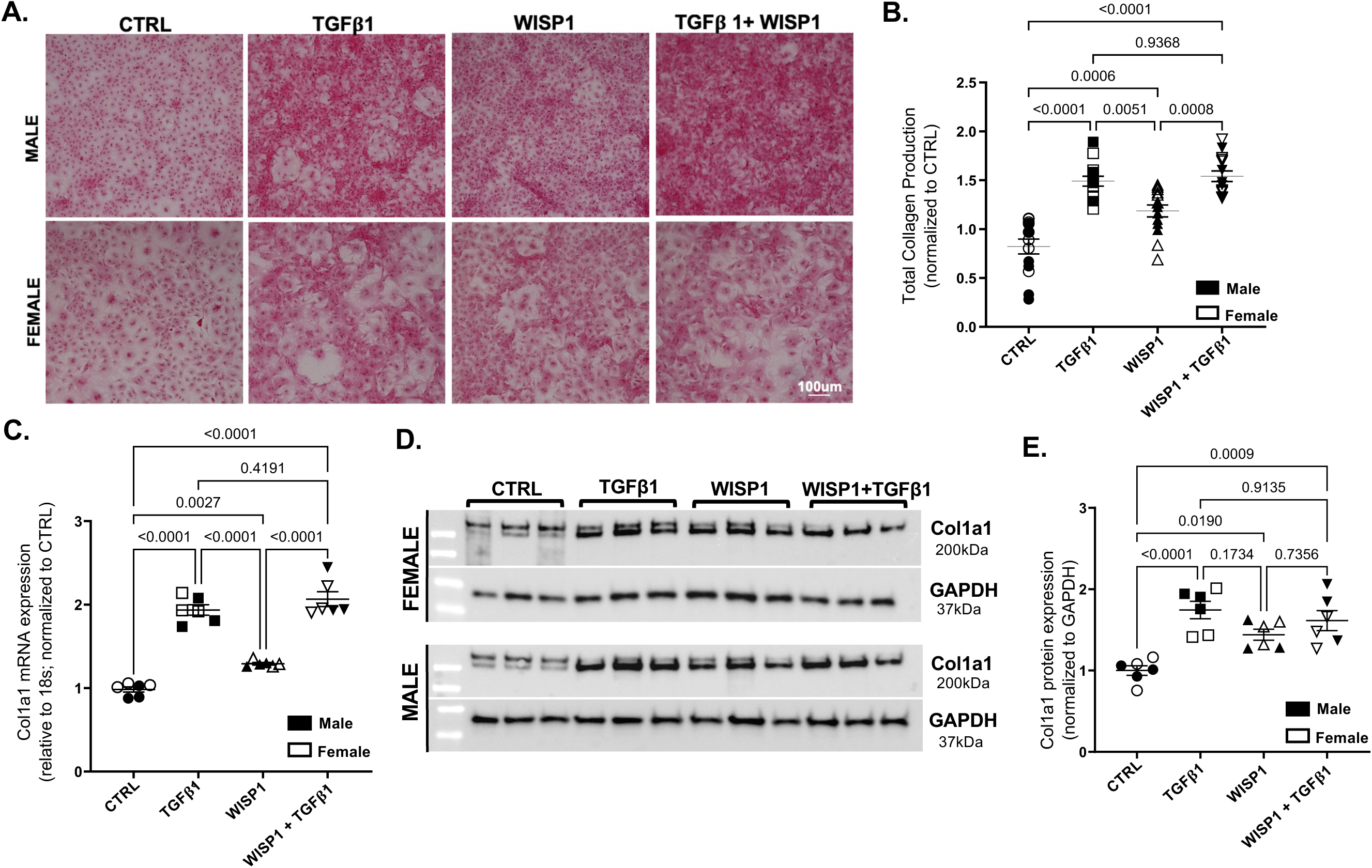
WISP1 enhances collagen expression and secretion in primary cardiac fibroblasts. (A–B) Total collagen production measured by picrosirius red elution (A540) following 72 h treatment (n = 14 isolations; 7 male, 7 female). **(C)** Col1a1 mRNA expression by qPCR. **(D–E)** Representative Western blot **(D)** and densitometric quantification **(E)** of collagen I normalized to GAPDH. Data are mean ± SEM. Statistical analysis by one-way ANOVA; *p < 0.05 versus Ctrl.

### 3.2. WISP1 independently induces fibroblast contractility and wound closure

We next evaluated the functional role of WISP1 on fibroblast contractile and wound healing activity using floating collagen gel contraction and scratch wound assays, respectively. In collagen gel contraction assays, WISP1 significantly enhanced gel contraction relative to control, though to a lesser extent than TGFβ1 (Fig. 3A-B). Similar to what was observed with α-SMA expression, results show that WISP1 further increases TGFβ1-dependent gel contractility (Fig. 3A-B). WISP1 also accelerated wound closure relative to control, but did not further enhance the TGFβ1 response (Fig. 3C-D). Together, the results thus far show that WISP1 independently induces myofibroblast marker expression and mediates functional ECM deposition, contractility, and wound healing, albeit to a slightly different extent with potential cooperativity with TGFβ1-mediated activity, hinting at differential mechanistic underpinnings.

**Figure 3.**
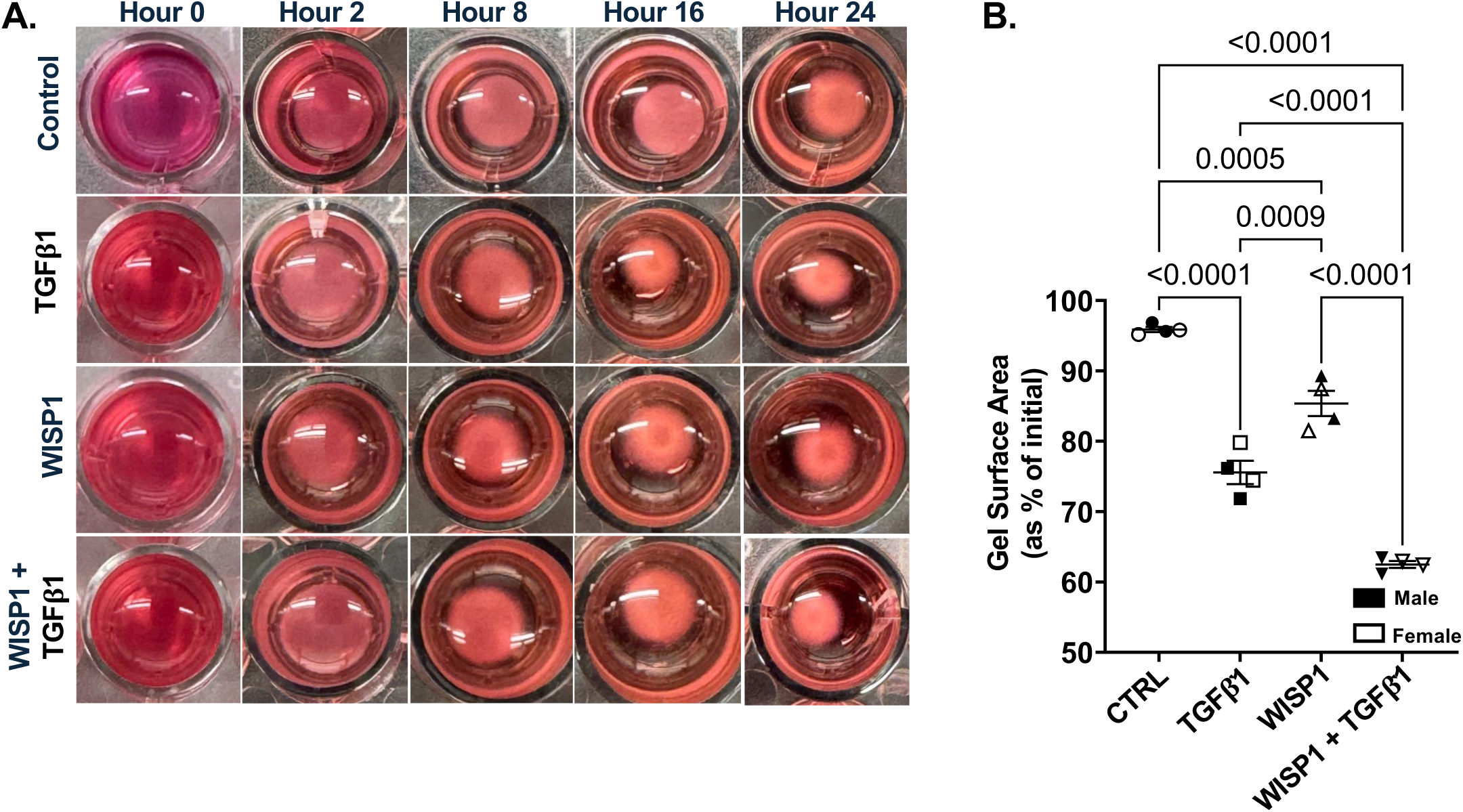
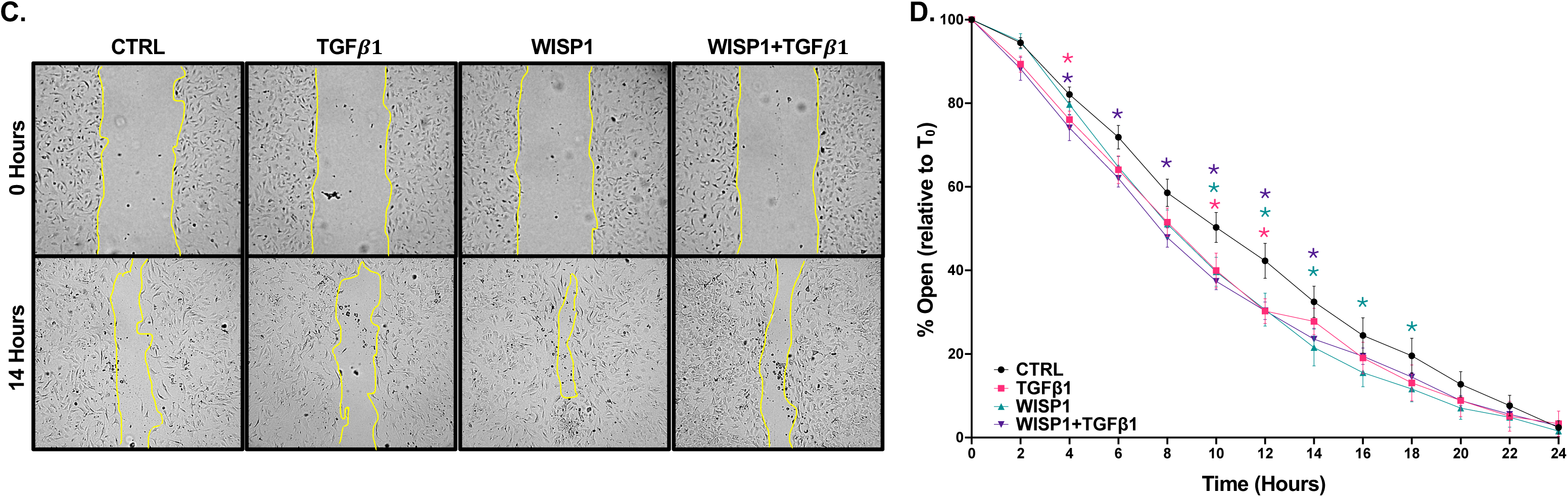
WISP1 enhances cardiac fibroblast contractility and accelerates wound closure (A–B) Primary adult mouse cardiac fibroblasts were embedded in floating collagen gels and treated for 24 h with vehicle (Ctrl), WISP1 (500 ng/mL), TGFβ1 (10 ng/mL), or WISP1 + TGFβ1. **(A)** Representative gel images at 0 h and 24 h. **(B)** Quantification of gel area (% contraction; n = 4 isolations, 2 males & 2 females). **(C–D)** Confluent fibroblast monolayers were scratched and imaged every 2 hours for 24 hours using the same treatment conditions. **(C)** Representative bright-field images at time point 0 and 14. **(D)** Quantification of wound closure (% of gap closed relative to time 0; n = 8 isolations, 4 male & 4 female). Data are mean ± SEM. One-way ANOVA with Tukey’s multiple comparisons; *p ≤ 0.05.

### 3.3. WISP1 mediates myofibroblast activity via a non-canonical signaling pathway

The data thus far suggested that WISP1 mediates many of the same myofibroblast pathways as TGFβ1. Thus, it was surprising to see that WISP1 failed to induce the expression of periostin (*Postn*), a canonical matricellular marker of myofibroblast activation, at either the RNA (Fig. 4A) or protein (Fig. 4B-C) level. To delineate the signaling pathways required for WISP1-mediated myofibroblast activity, we co-treated with SB203580, a selective inhibitor of p38 MAPK, an established non-canonical pathway of myofibroblast activation. Western blot showed that p38 inhibition significantly blunted the WISP1-dependent expression of α-SMA, but did not alter TGFβ1-dependent expression (Fig. 4D-E). Similarly, p38 inhibition did not affect the TGFβ1-dependent expression of periostin, which we showed to be independent of WISP1 (Fig. 4F-G). Finally, picrosirius red quantification of decellularized ECM shows that p38 inhibition also completely abrogated WISP1-mediated total collagen deposition (Fig. 4H). Interestingly, in the case of α-SMA and total collagen, but not periostin, p38 inhibition also resulted in a slight reduction in the baseline and WISP1+TGFβ1 levels, suggesting a WISP1-p38 signaling dependency in both of these groups. Together, this data shows a divergence of WISP1-mediated myofibroblast activity from canonical TGFβ1 signaling and suggests, by lack of periostin expression, a unique myofibroblast cell state following WISP1 activation.

**Figure 4.**
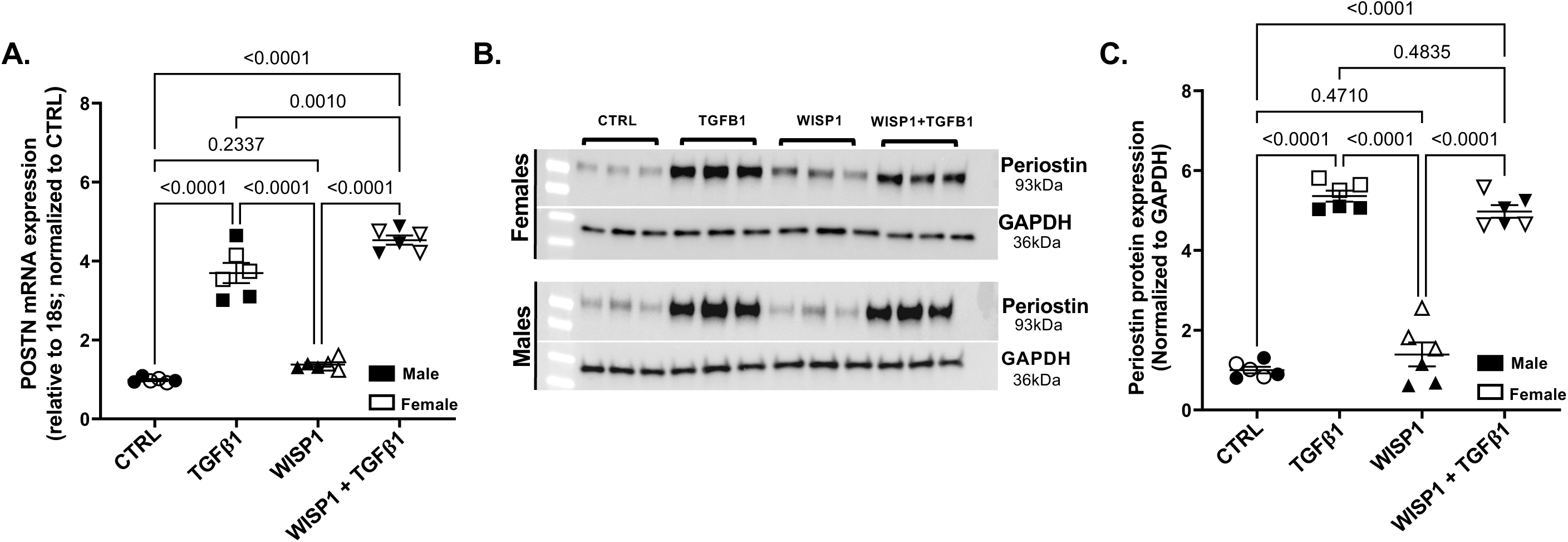
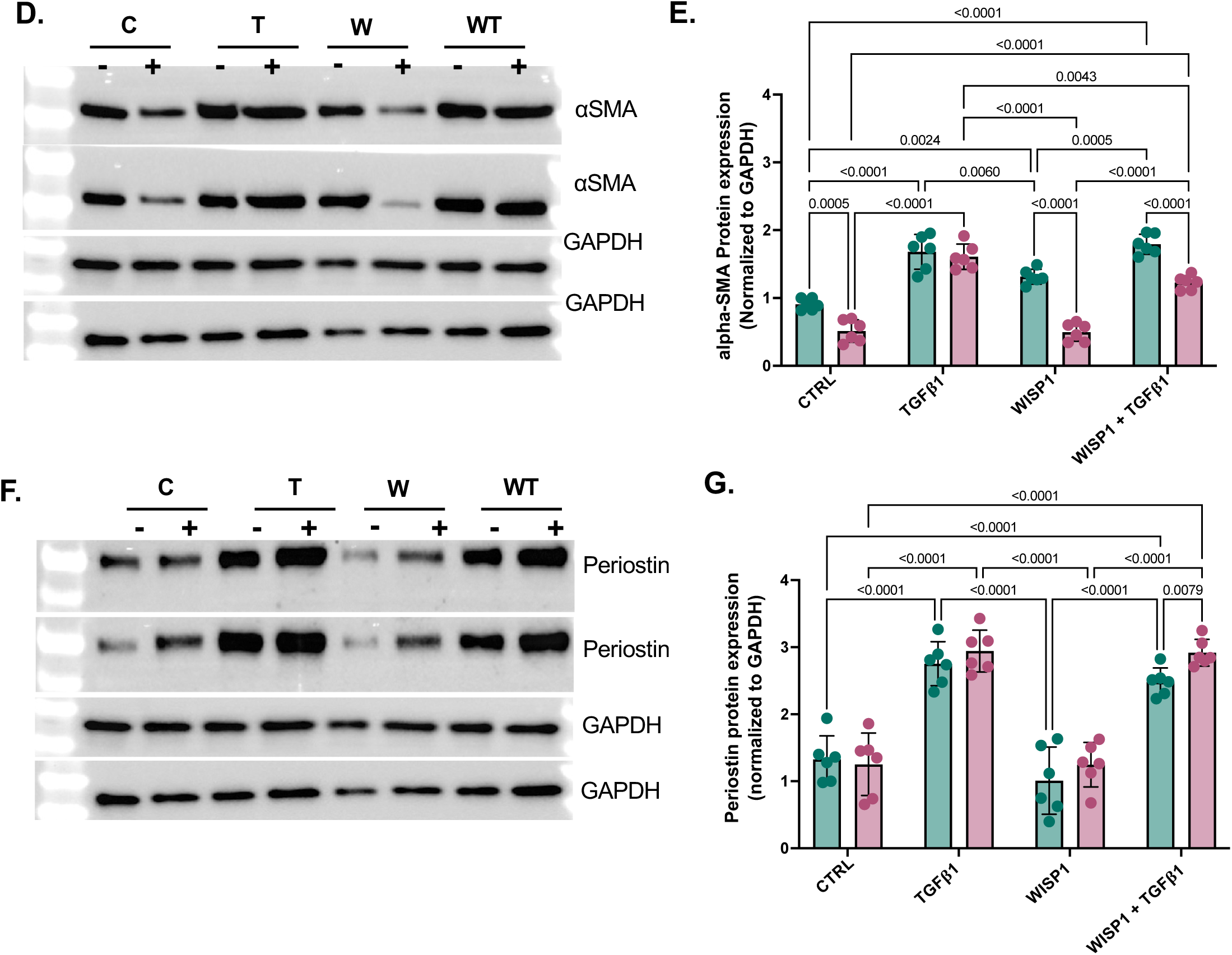
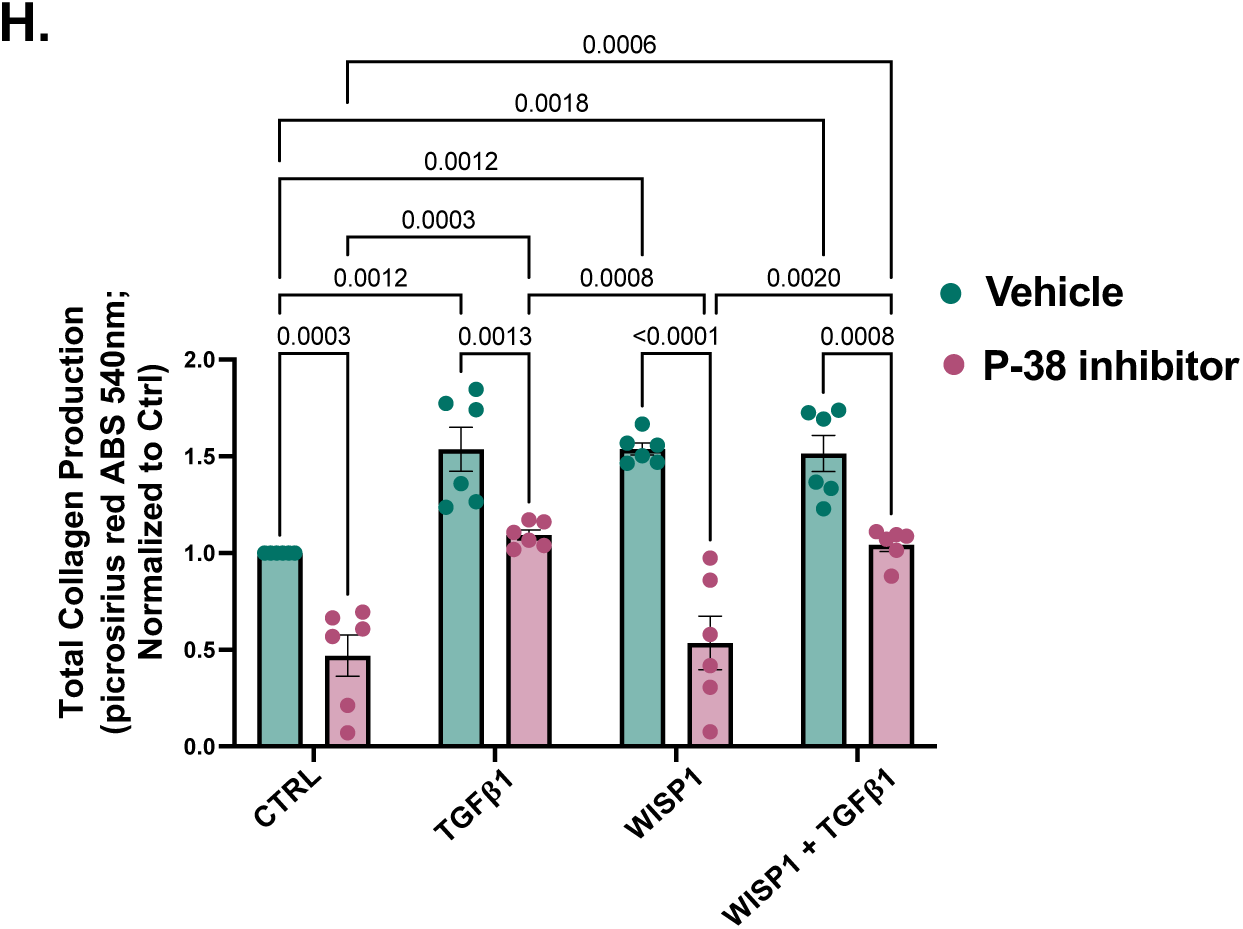
WISP1-dependent myofibroblast activation is p38 MAPK–dependent and uncoupled from periostin expression. **(A)** Periostin (*Postn*) expression assessed by qPCR in primary cardiac fibroblasts treated for 72 h. **(B-C)** Representative Western blot **(B)** and densitometry normalized to GAPDH and Ctrl **(C)** of Periostin across 6 biological replicates. **(D-E)** Representative Western blots for α-SMA **(D)** and densitometric quantification **(E)** following treatment with p38 MAPK inhibitor SB203580 (10 μM). **(F-G)** Representative Western blots for periosotin **(F)** and densitometric quantification **(G)** following treatment with p38 MAPK inhibitor SB203580 (10 μM). **(H)** Total soluble collagen secretion measured by Picrosirius Red. Data are mean ± SEM. Statistical significance was determined by one-way ANOVA or two-way ANOVA as appropriate.

### 3.4. The Wisp1-dependent transcriptome is enriched for gene expression associated with proliferative and inflammatory signaling

We next took an unbiased multi-omics approach, to fully characterize the WISP1-dependent changes within the cardiac fibroblast transcriptome and proteome. We first assessed transcriptomic changes via bulk RNA-sequencing in cardiac fibroblasts treated with TGFβ1, WISP1, or WISP1 + TGFβ1 for 72 hours. In order to capture broad WISP1-dependent changes across the transcriptome that were not limited by arbitrary thresholds, we performed rank-based gene set enrichment analysis (GSEA) to identify pathway-level WISP1-dependent gene expression networks.

GSEA results show that Wisp1 induces a unique transcriptomic signature compared to TGFβ1, with a WISP1-specific increase in proliferation (Fig. 5A-B) and inflammation (Fig. 5A, C) associated pathways. Consistent with these pathway-level patterns, mean nodule enrichment scores (NES) demonstrated a net positive shift in proliferation and immune/inflammation signatures following WISP1 treatment (Fig. 5D). In contrast, TGFβ1 produced a transcriptional state characterized by suppression of proliferative and inflammatory pathways, while enhancing canonical stress and fibrosis associated gene programs (Fig. 5A, D). Notably, whereas TGFβ1 robustly increased canonical ECM and myofibroblast module scores, WISP1 alone showed minimal induction of these modules (Fig. 5A, E), supporting a WISP1-driven transcriptome that is not dominated by canonical profibrotic differentiation programs.

**Figure 5.**
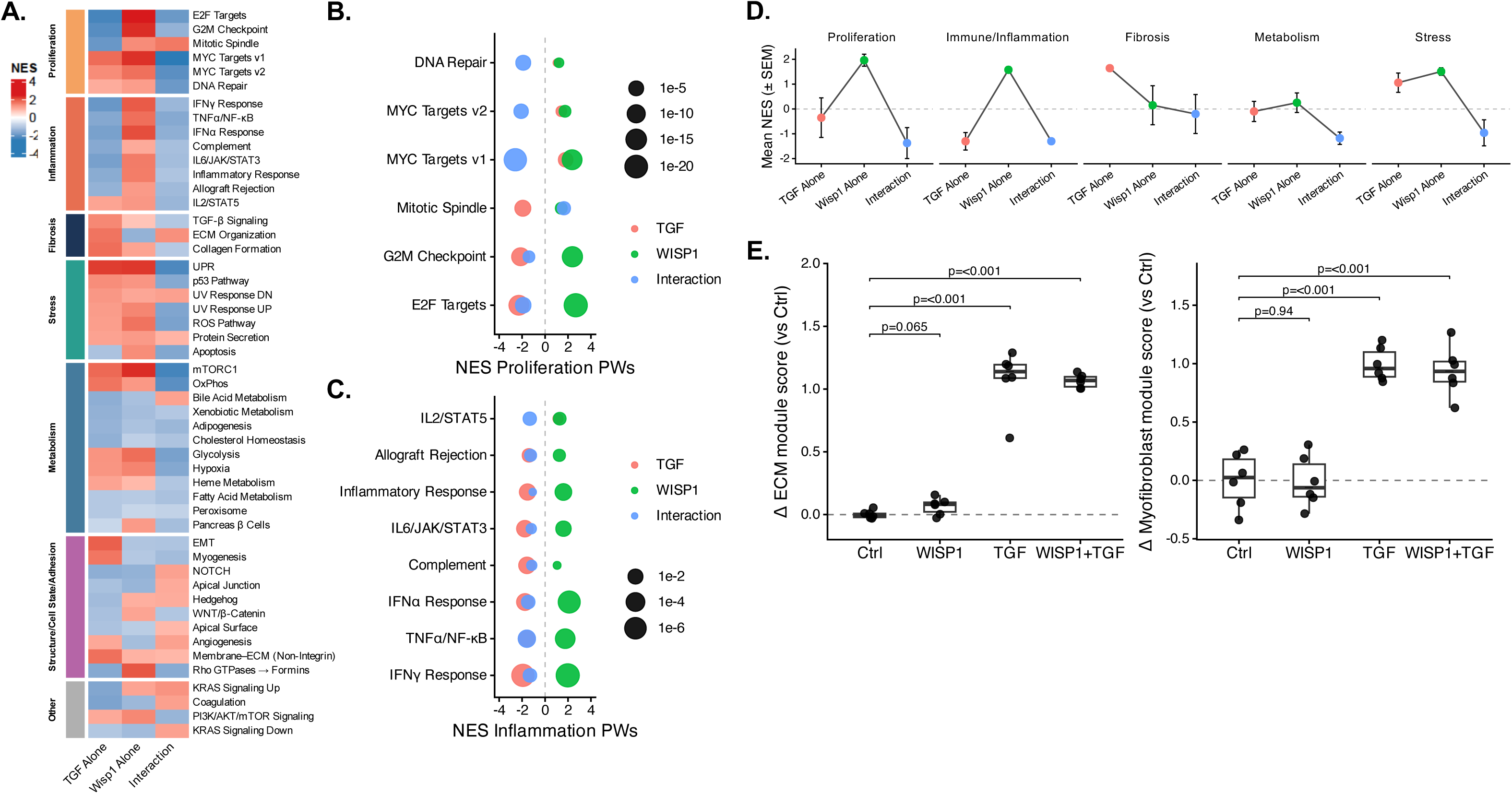
WISP1 establishes a proliferative and immune-inflammatory transcriptomic program distinct from TGFβ1-driven myofibroblasts. **(A)** Heatmap of Hallmark gene set normalized enrichment scores (NES) across treatments. **(B–C)** Dot plot representation of proliferation **(B)** and inflammation **(C)** pathway enrichment. **(D)** Aggregated mean NES across functional categories. **(E)** Myofibroblast and ECM module scores relative to control. Data presented as mean ± SEM or box-and-whisker plots.

The combined WISP1+TGFβ1 treatment did not produce additive activation of WISP1 or TGFβ1-driven gene expression. Instead, the interaction revealed a widespread antagonism of the WISP1 proliferation and inflammation gene programs (Fig. 5B, D), consistent with a context dependent fibroblast activation patter by both WISP1 and TGFβ1. At the module level, TGFβ1 and WISP1+TGFβ1 produced similarly elevated ECM and myofibroblast module scores (Fig. 5E), indicating that WISP1 does not further increase the overall magnitude of the canonical TGFβ1-driven myofibroblast transcriptional program. A heatmap of individual genes used to extrapolate the myofibroblast and ECM module scores can be found in Fig. S2.

Together, these data identify WISP1 as a context-dependent regulator of fibroblast transcriptomic state that promotes proliferative and immune–inflammatory gene expression networks while failing to robustly induce canonical ECM/myofibroblast transcriptional changes.

### 3.5. WISP1 drives cell compartment-specific changes in the proliferative and inflammatory proteome

To determine whether the WISP1-dependent transcriptional state translated into functional protein-level remodeling, we performed label-free quantitative mass spectrometry on the global soluble (intracellular cytosolic and nuclear) fraction as well as the decellularized ECM fraction (Fig. S3), and used these datasets to further extrapolate an ECM-enriched subset defined by proteins preferentially detected in the ECM relative to the soluble fraction. GSEA was again applied to ranked protein abundance changes to quantify coordinated pathway-level responses across treatments (Fig. 6A).

**Figure 6.**
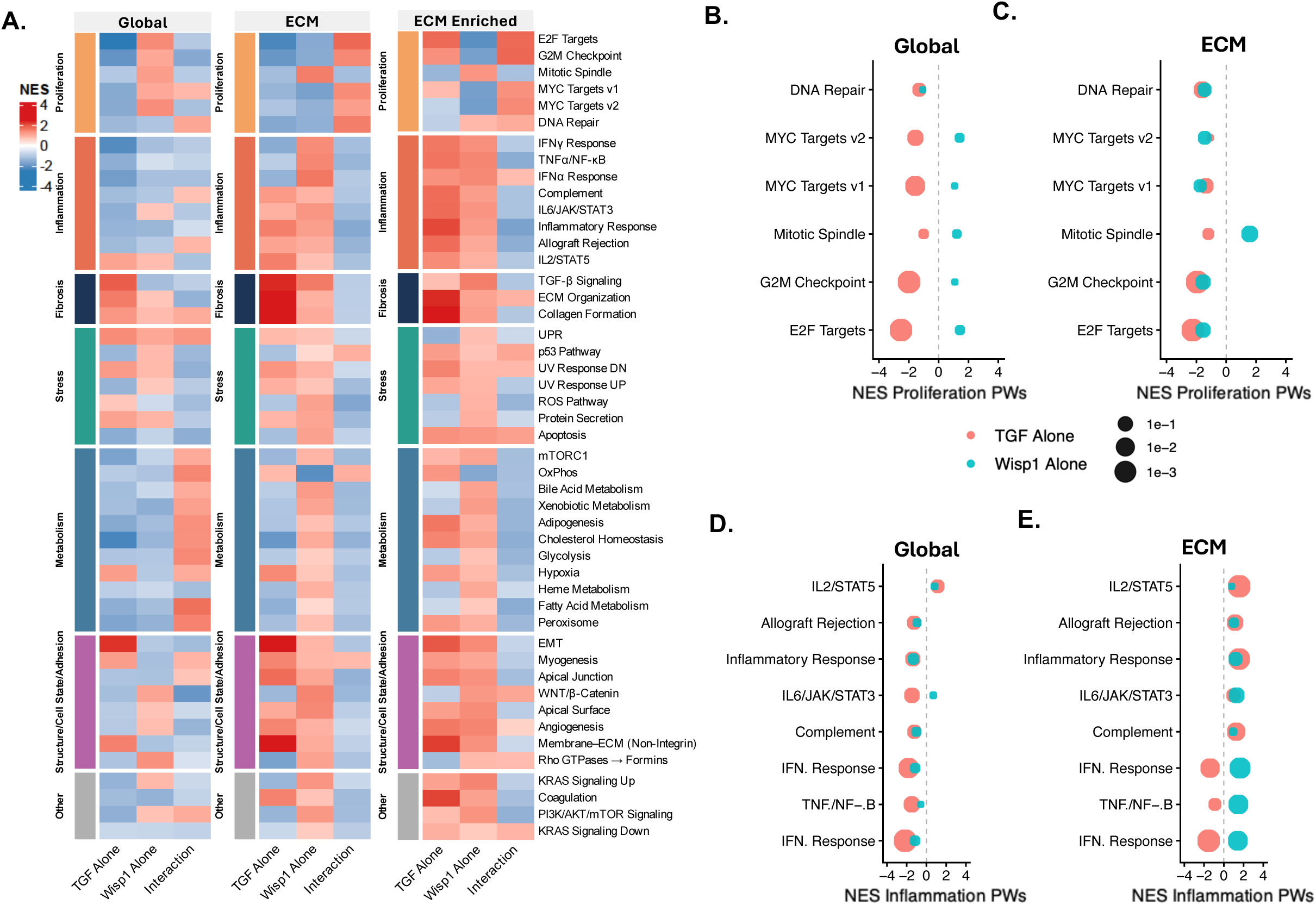
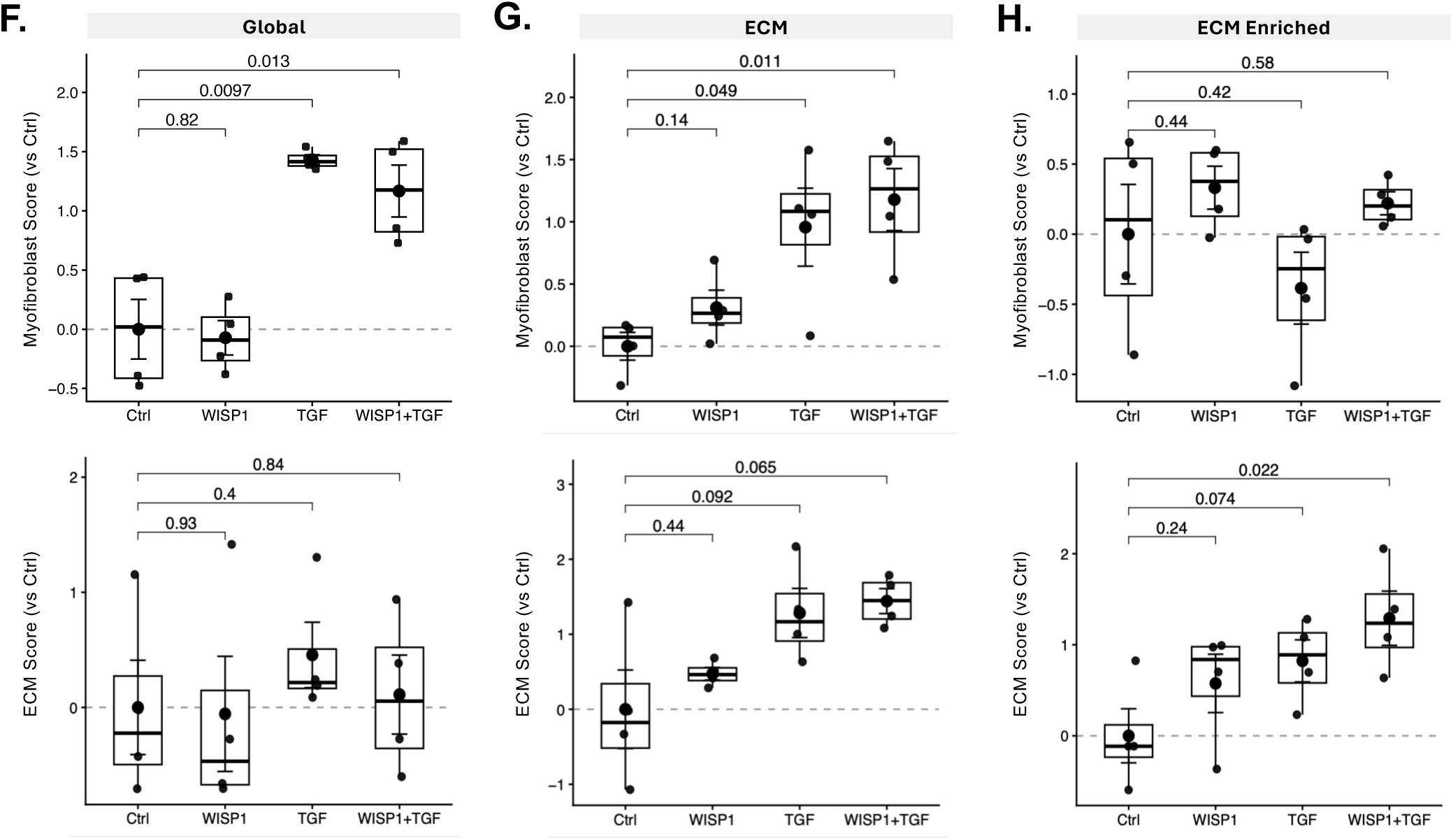
Compartment-resolved proteomics reveals WISP1-driven intracellular proliferation and ECM-associated immune remodeling. **(A)** Heatmap of pathway NES across global (soluble), ECM, and ECM-enriched proteomes. **(B–C)** Dot plot analysis of proliferation and inflammation pathway enrichment in global **(B)** and ECM **(C)** fractions. Dot size represents significance as adjusted p-value. **(D–F)** Myofibroblast and ECM module scores across global **(D)**, ECM **(E)**, and ECM-enriched **(F)** proteomes. Box plots show median with interquartile range.

In the global intracellular proteome, WISP1 drives a proteomic signature consistent with the increase in a proliferative transcriptomic program observed in RNA-seq (Fig. 6A–B). In parallel with the RNA-seq results, TGFβ1 alone exhibited negative enrichment across proliferation associated protein groups (Fig. 6A-B). Similarly, WISP1-mediated an increase in inflammatory programs, with an inverse TGFβ1-mediated suppression of the same pathways, but the inflammatory proteome was enriched in the ECM fraction (Fig. 6A, D-E). This is in contrast to the WISP1-dependent proliferation-associated proteome, which was enriched in the global/intracellular fraction (Fig. 6A-C). In addition to an antagonism of these WISP1-dependent protein networks, TGFβ1 expectedly promotes the expression of canonical fibrosis-associated proteins, with an enrichment of these proteins in the ECM compartment (Fig. 6A). The compartmentalized regulation of these pathways between the intracellular and ECM proteome is expected given signaling nature of each with proliferation-associated proteins acting intracellularly, and fibrosis and inflammation-associated proteins being predominately secreted into the ECM compartment.

Similar compartmentalization is observed in the myofibroblast and ECM module scores, where TGFβ1 induces a significant enrichment in myofibroblast score in the global and ECM proteomes, but only yields a significant increase in the ECM module score within the ECM proteome fraction (Fig. 6F-G). In accordance with the RNA-seq data, WISP1 does not significantly increase either the myofibroblast or ECM score in either proteome fraction (Fig. 6F-G; a heatmap of individual proteins used to extrapolate the myofibroblast and ECM module scores can be found in Fig. S4). As expected, the ECM enriched proteome shows a stronger trend to increased ECM related protein module scores in all cases compared to myofibroblast proteins modules (Fig. 6H and Fig. S3).

Taken together, the proteomics results demonstrate that WISP1 promotes a proliferative and immune modulatory state with compartmentalized enrichment of these proteins in the expected intracellular and ECM fractions, respectively, while TGFβ1 suppresses these pathways and induces canonical myofibroblast protein expression.

## 4. Discussion

Cardiac fibroblast activation is frequently thought of as a largely linear progression toward a canonical, TGFβ-driven myofibroblast phenotype that primarily promotes extracellular matrix remodeling. WISP1 has been reported to increase in the heart after myocardial infarction and has been suggested to play a role in fibrotic remodeling through myofibroblast activation [27, 30, 31, 32]. Here, we identify WISP1 as a context-dependent regulator of cardiac fibroblast activation that elicits some of the hallmark features of myofibroblasts including α-SMA expression, collagen production, contractility, and wound closure [33]. However, WISP1 fails to induce the expression of periostin, a primary myofibroblast marker gene [34]. Instead, WISP1 produces a myofibroblast cell state that is distinct from a TGFβ1-driven classical myofibroblast. Our data support WISP1 promoting a mechanically active fibroblast with only a weak induction of traditional ECM remodeling pathways but instead possesses a transcriptome and proteome that is instead primarily enriched for proliferation and immune.

Our results showing WISP1-dependent α-SMA and collagen expression corroborates previous literature in a more robust and quantitative fashion [27,28]. Following up on this data, our demonstration of WISP1-induced contractility and wound healing also fits a model of WISP promoting a canonical myofibroblast phenotype. However, our finding that WISP1 fails to induce canonical myofibroblast expression of periostin was unexpected as periostin is widely used as a marker of myofibroblast differentiation and is implicated in ECM organization and stabilization during cardiac remodeling [35, 36]. This result suggested that WISP1-dependent remodeling alters ECM composition or organization in ways that differ significantly from canonical profibrotic remodeling.

Mechanistically, we further show that WISP1-driven activation depends on non-canonical p38 MAPK signaling, providing a molecular signaling basis for its divergence from TGFβ1-driven myofibroblast activity. Pharmacologic p38 inhibition abrogated WISP1-induced α-SMA expression and collagen deposition, while having minimal impact on their TGFβ1-driven induction. Importantly, periostin remained TGFβ1-responsive and unaffected by p38 inhibition, consistent with a model that periostin is regulated through canonical profibrotic signaling [34]. This result is also consistent with recent work by Li et al that suggested WISP1 mediates α-SMA and collagen processing downstream of its binding to the α5β1 integrins [28].

Our transcriptome and proteome multiomics results provide a much deeper probe into WISP1-dependent mechanisms that had not yet been done. The primary novel finding from these concordant datasets is that the dominant molecular signature of WISP1 is not a classical ECM program, but a proliferative and immune–modulatory axis that is respectively compartmentalized across the intracellular and ECM proteomes. Transcriptomic pathway analysis revealed strong WISP1-dependent enrichment of proliferation and inflammatory signaling programs, whereas TGFβ1 robustly suppressed these networks while inducing canonical pro-fibrotic and ECM remodeling gene networks. Surprisingly, the WISP1 and TGFβ1 co-treatment group did not reflect additive activation, but rather indicated an antagonism of WISP1-driven gene expression programs, suggesting that WISP1 and TGFβ1 promote opposing transcriptional states. These results demonstrate that WISP1 promotes a unique myofibroblast state and it not a profibrotic agonist as previously postulated.

The cell compartment level resolution in our proteomics data further shows that WISP1’s proliferative program is most prominent in the global (cytosolic/nuclear) proteome, whereas the enrichment in inflammatory proteins occurs in the ECM compartment. This divergence is consistent with expected cellular organization in that proliferation networks are predominantly intracellular, while immunomodulatory proteins are secreted or matrix-associated. Our data shows consistency of these WISP1-dependent proteins with specific compartmentation while remaining a comparatively weak driver of canonical ECM/myofibroblast protein modules, both of which are consistent with our RNA-seq results. In contrast, TGFβ1 increased myofibroblast and ECM module scores most robustly in the ECM-associated fractions, consistent with classical profibrotic remodeling. Importantly, the ECM-enriched subset emphasized that the major influence of WISP1 is not simply to raise bulk secretion to the ECM, but appears specific to inflammation-associated proteins. This compartmentalization supports a model in which WISP1 may promote the transition to a myofibroblast that contributes to the myocardial remodeling environment through both mechanical activity with strong implications immune-modulatory cell–cell signaling in the injured myocardium.

Our findings are consistent with, but also extend, prior work implicating WISP1 in fibrotic and remodeling contexts. WISP1 expression is low in the healthy adult heart and increases following injury, and previous studies in other organs have linked increased WISP1 expression to fibroblast proliferation, migration, and ECM synthesis. However, those models position WISP1 primarily downstream of TGFβ1 signaling as a profibrotic effector. By explicitly contrasting WISP1 and TGFβ1 in the same cells and integrating functional assays with transcriptomics and proteomics, we show that WISP1 drives a specific myofibroblast phenotype that diverges from classical TGFβ1-driven fibrosis. This distinction may help reconcile why WISP1 has been associated with fibrotic remodeling in vivo, yet does not functionally recapitulate the canonical profibrotic myofibroblast phenotype.

A noted limitation to our results is the necessity to perform these experiments in cultured primary adult mouse cardiac fibroblasts while acknowledging that in vivo fibroblasts are additionally influenced by dynamic paracrine signals, mechanical load, and cell-cell interactions. Additionally, our ECM proteomics data provides a practical compartmental view of extracellular remodeling, it does not directly measure functional ECM parameter such as extent of crosslinking or biomechanical properties. Of note, these are also parameters likely to be impacted by the presence of lack of periostin expression. Finally, our data support context-dependence between WISP1 and TGFβ1, but the mechanistic basis of this interaction antagonism or the relative temporal signaling between WISP1 and TGFβ1 remains to be defined.

It also remains unclear how temporal or spatial expression of WISP1 and TGFβ1 in vivo coordinate to give rise to distinct myofibroblast cell states, and a better understanding of this will be important for translating these results into therapeutic application. Cell type-specific genetic perturbation of Wisp1 (and/or its receptors) is needed to define whether WISP1 primarily contributes to early maladaptive ECM remodeling, immune recruitment, or scar maturation/myofibroblast resolution in translationally relevant injury models. Finally, given the WISP1-dependent expression of immune/inflammatory gene and protein networks, it will be critical to determine the functional role that WISP1 plays in mediating fibroblast to immune cell crosstalk, and whether it contributes to myeloid cell recruitment, macrophage inflammatory polarization, or inflammatory resolution during myocardial remodeling.

In summary, our data indicate that WISP1 promotes collagen production and contractile behavior in cardiac fibroblasts, but does not promote the transcriptional or proteomic changes indicative of a classical TGFβ1-driven myofibroblast activation. Instead, this work repositions WISP1 from a putative pro-fibrotic effector to an immune modulatory state regulator of myofibroblasts and provides a mechanistic basis for how matricellular proteins such as WISP1 contribute to diversity among fibroblast phenotypes during cardiac injury and repair. In conclusion, our integrated transcriptomic and spatially compartmentalized proteomic results show that WISP1 acts as a context-dependent mediator of a novel and unique novel cardiac myofibroblast cell state that couples mechanical activation with an immune-modulatory phenotype rather than canonical fibrotic matrix remodeling.

## 5. Data Availability Statement

All data will be publicly available following peer-review.

## 6. Funding

This work was supported by NIH grants R01-HL166326 (MT and OK), R01-HL148598 (OK), and American Heart Association Transformational Project Award 24TPA1290119 (MT). SP was supported by American Heart Association Predoctoral Fellowship Award 1029875, NIH Predoctoral Fellowship F31-HL170636 and NIH Training Grant T32 HL125204.

## Acknowledgements

We acknowledge resources from the Campus Chemical Instrumentation Center Mass Spectrometry and Proteomics Facility and the OSU Comprehensive Cancer Center (OSUCCC) Proteomics Shared Resource (PSR), and the Genomics Shared Resource Center at The Ohio State University. The Cytek Aurora flow cytometer was made available by The Ohio State University Comprehensive Cancer Center (OSUCCC) and the National Institutes of Health (NIH) under grant P30CA016058. This research was made possible through resources, expertise, and support provided by the Pelotonia Institute for Immuno-Oncology (PIIO) and the Immune Monitoring and Discovery Platform, which is funded by the Pelotonia community and the OSUCCC. The content is solely the responsibility of the authors and does not necessarily represent the official views of the National Institutes of Health.

## Notes

### Competing Interest Statement

The authors have declared no competing interest.

